# The receptor VLDLR binds East Equine Encephalitis virus through multiple distinct modes

**DOI:** 10.1101/2023.11.30.569340

**Authors:** Duanfang Cao, Bingting Ma, Ziyi Cao, Xiaoyu Xu, Xinzheng Zhang, Ye Xiang

## Abstract

Eastern Equine Encephalitis virus (EEEV) is an alphavirus that can cause severe diseases in infected humans. The very low-density lipoprotein receptor (VLDLR) was recently identified as a receptor of EEEV. Herein, we performed cryo-electron microscopy structural and biochemistry studies on the specific interactions between EEEV and VLDLR. Our results show that VLDLR binds EEEV at three different sites A, B and C through its membrane-distal 8 LDLR class A (LA) repeats (LA1-8). Site A is located in the cleft in between the E1-E2 heterodimers of the surface trimeric spikes. Site B is located near the connecting β ribbon in between the B and A-C domains of E2 and is in proximity to site A, while site C is on the domain B of E2. LA1 and LA5 specifically recognize site A, whereas LA2, LA3 and LA6 can bind both sites A and C. The binding of VLDLR LAs to EEEV is in complex modes. The LA1-2 mediated binding plays a critical role in mediating virus entry. The mutation W132G of VLDLR impairs the binding of LA3, drives the switch of the binding modes, and significantly enhances the attachment of EEEV to the cell. The W132G variant of VLDLR could be identified in human genome and SNP sequences, implying that people with similar mutations in VLDLR may be highly susceptible to EEEV infection.

---

The Eastern equine encephalitis virus (EEEV) is an alphavirus that can cause severe and sometimes fatal disease in humans and equines^1,2^. EEEV is commonly found in the eastern regions of America and has been associated with outbreaks of encephalitis^3^. Birds are the major enzootic hosts and nature reservoir of EEEV. EEEV is transmitted among birds mainly through the mosquito *Culiseta melanura*^4^. Other mosquito species, like *Aedes* and *Coquillettidia*, can also transmit EEEV and are believed to be the bridge vectors for the transmission of EEEV from birds to humans and horses^4,5^. Unlike the encephalitic alphavirus Venezuelan equine encephalitis virus (VEEV) that efficiently amplify and develop a high viremia in infected equines, EEEV infection of humans or horses can not develop a high viremia that is sufficient for establishing a complete transmission cycle^6,7^. Antigenic variants of EEEV are categorized into two major groups, including the more virulent North American variant and the less virulent South American Variant^8^. So far, there is still no available vaccine or drug for prevention and treatment of EEEV infection in humans.

The spherical virions of EEEV are approximately 70 nm in diameter and are assembled by the viral RNA genome, the envelope proteins E1, E2, the capsid protein C and the small 6 kDa protein 6K^9–11^. The envelope and capsid proteins of EEEV assemble to form two concentric protein shells that are separated by a lipid bilayer. The envelope protein shell is composed of 80 trimeric spikes that are arranged with T=4 icosahedral symmetry^12–15^. Twenty of the 80 spikes are sitting on the icosahedral 3 fold axes (i3) and sixty of the 80 spike are sitting on the quasi 3 fold axes (q3). Each trimeric spike is an assembly of three E1 and E2 heterodimers and can be divided into two distinct parts, including a flower-like protruding portion and a shell-forming portion. The flower-like protruding portion is composed by the three immunoglobulin-like β domains A (residues 1-132), B (residues 173-231), and C (residues 269-342) of E2 and the immunoglobulin-like β domain 2 (residues 40-127 and 170-272) of E1 (E1- DII)^10,11,15^. Domains A and C of E2s are located in the center of the spike and are surrounded by the E1-DIIs. The three E2 domain Bs are arranged anti-clockwisely on the top of the spike and constitute three petals of the flower. The B domain of E2 is connected to the domains A and C of E2 through two long linker peptide (residues 133- 172, 231-268) named the “β-ribbon connector”. The E1-DI (residues 1-36, 133-168, 274-293) and DIII (residues 294-381) at the base of the spike splay tangentially along the viral surface and interact with each other to constitute a smooth protein shell^15^.

Like other alphaviruses, EEEV enters host cells through receptor-mediated endocytosis. The spikes of the E1-E2 heterodimers are responsible for specific receptor attachment^10,11^. Recently, the very low-density lipoprotein receptor (VLDLR) and the apolipoprotein E receptor 2 (ApoER2), two closely related receptors from the low-density lipoprotein (LDL) receptor family, were identified as the receptors of EEEV^16^. LDLRAD3 of the same LDL receptor family was identified as an entry receptor of VEEV^17^. VLDLR and ApoER2 can also mediate the entry of Semliki Forest virus (SFV) and Sindbis virus (SINV), two alphaviruses that rarely cause encephalitis in equines or humans^16^. The Matrix Remodeling Associated protein 8 (MXRA8) was identified as an entry receptor of many other alphaviruses^18,19^.

Receptors from the LDL receptor family have similar small repeats in their ectodomains^20^. The small repeat domain has approximately 40 amino acid residues and a compact fold that contains a calcium binding motif and is crosslinked by three disulfide bonds^21–23^. Although both VLDLR and LDLRAD3 bind the viruses through their ecto small repeat domains, these two receptors bind the viruses with distinct different modes. LDLRAD3 binds through its distal repeat domain in clefts between the E1-E2 heterodimers of the protruding VEEV spikes^24,25^. However, VLDLR binds SFV on the smooth shell constituted by E1-DIIIs in a synergistic mode that involves multiple ecto repeat domains of the receptor^26^. Comparing to VLDLR and LDLRAD3, MXRA8 has a quite different structure and binds the viruses through its distal Ig-like domains. Of note, MXRA8 bind in clefts between the E1-E2 heterodimers of the protruding spikes as does LDLRAD3, although the interactions are completely different^18,27,28^. Our recent results showed that it is unlikely that VLDLR binds EEEV in a similar mode as does SFV. The structural basis for the selective binding of VLDLR to EEEV need to be further investigated.

## Structure of VLDLR LA1-8 in complex with EEEV

The membrane-distal homologous LDLR class A (LA) repeats of VLDLR and ApoER2 were shown to mediate the interactions between EEEV and VLDLR^16^. ApoER2 have two isoforms and the isoform2 (ApoER2iso2) is the prodominent form. VLDLR contains eight LA repeats (VLDLR LA1-8), while ApoER2iso2 contains three LA repeats (ApoER2iso2 LA1-3). Sequence alignments show that the sequence identity between LA1-3 of VLDLR and LA1-3 of ApoER2iso2 is 55.4% (Extended Data Fig. 1a). We, thus, produced VLDLR LA1-8 and ApoER2iso2 LA1-3 in fused with an antibody constant region fragment at the C-terminus, and prepared the complexes with the EEEV virus-like particles (VLPs), which were produced by using the structural proteins of the EEEV PE6 strain^29^. The structures of the complexes were determined by using cryo-electron microscopy (cryo-EM) single particle reconstruction at a resolution of 5.3 Å for the VLDLR LA1-8-Fc and EEEV PE6 VLP complex and 4.9 Å for the ApoER2iso2 LA1-3 and EEEV PE6 VLP complex with icosahedral symmetry imposed (Extended Data Fig. 1b-e, Extended Data Table 1). Compared to the LA free EEEV PE6 VLPs, we identified additional densities on the spikes of VLDLR LA1-8-Fc-EEEV PE6. However, we did not observe additional density in the reconstruction of ApoER2iso2 LA1-3-EEEV PE6 (Extended Data Fig. 2a). Further block-based reconstructions with the asymmetric unit as a block were calculated for the VLDLR LA1-8-Fc and EEEV PE6 VLP complex and yield a density map at a resolution of 3.5 Å (Extended Data Fig. 1b-d).

Additional densities at three distinct sites A, B and C were observed on the spikes of VLDLR LA1-8-Fc-EEEV PE6. Site A is located in a cleft formed in between two neighboring E1-E2 heterodimers (Fig. 1a-c), a similar location where LDLRAD3 binds to VEEV^24,25^. Site B is located near the connecting β strands in between the A and B domains of the EEEV PE6 E2 protein (Fig. 1a-c). Site B is adjacent to site A. Densities at site B are much weaker compared with these at site A. Weak densities were observed in between sites B and A at a low contouring level (Fig. 1b). Site C is on the B domain of E2 and is approximately 60 Å and 40 Å away from sites A and B, respectively (Fig. 1a-c). The densities on the three binding sites are good enough for model fitting but not for ab initio model building of the bound VLDLR LAs (Extended Data Fig. 2b). We, thus, were not able to unambiguously distinguish the bound LAs from the 8 LA repeats. Of note, densities at different A sites in the asymmetric unit show distinct different features (Fig. 1d).

**Fig. 1.**
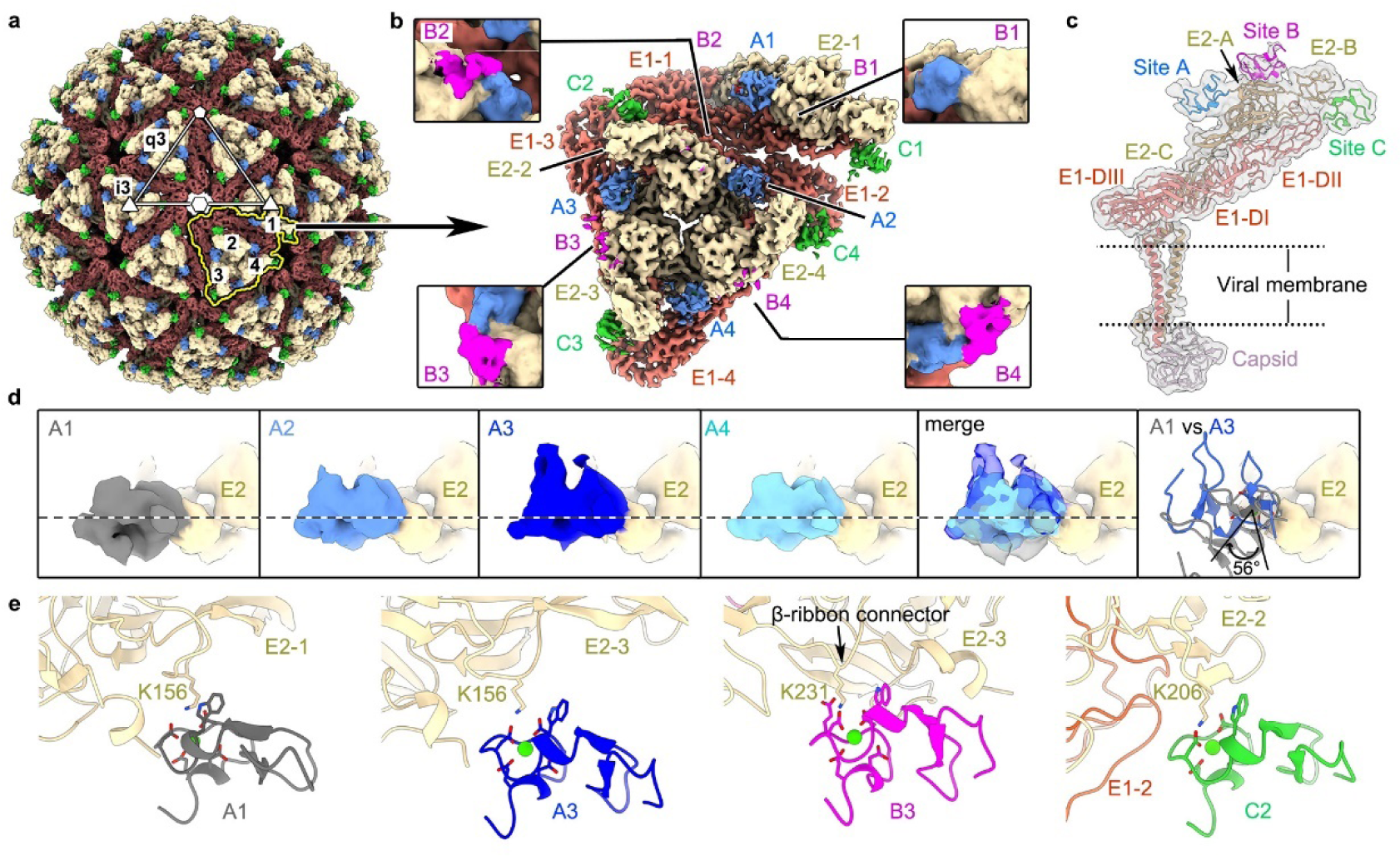
S**t**ructure **of VLDLR LA1-8 in complex with EEEV PE6 VLP. a**, A surface rendered representation showing the density map of EEEV PE6 VLP bound with LA1-8-Fc. The E1, E2, and capsid of the EEEV PE6 VLP are colored coral, wheat, and thistle, respectively. Additional densities at two different locations on the spike, which could be contributed by the bound LA repeats, are colored blue and green, respectively. The white triangle indicated one asymmetric unit of the E protein shell. The outlines of four E1-E2 heterodimers in one asymmetric unit are shown in yellow. The labels i3 and q3 indicate one i3 spike and one q3 spike, respectively. **b**, A surface rendered representation showing the density map of one asymmetric unit. The density map was obtained from the block-based reconstruction that treats one asymmetric unit as an individual block. The E proteins are colored the same as in “a”. The additional densities at three different sites on the spike are colored blue, green and magenta, respectively. The zoom-in inserts show the densities at sites B. The density map in the zoom-in inserts was low pass filtered to a resolution of 10 Å and the contouring level was set to 0.0035 e^-^/Å^3^. **c**, A cut-off view of the spike showing the positions of E proteins, capsid and the bound LA repeat at sites A, B and C. The density map is set transparent. The fitted structures are shown in ribbon with E1, E2 and capsid proteins colored the same as in “a”. Fitting of the LA3 structure (pdb accession number: 8IHP) into the cryo-EM density showing the orientation and position of the bound LA repeats, which are colored blue, magenta and green, respectively. **d**, Comparison of the LA densities at the four A sites. **e**, Ribbon diagrams show the interactions between the E protein and the bound LAs. The potential key residues involved in the interactions are shown in sticks. The calcium ions of the bound LAs are shown in green balls.

Structure predictions of the LAs^26,30^ showed that they have a similar structure with an elliptic cylinder shape (Extended Data Fig. 2c). The LA has two ridges and two flat surfaces (Extended Data Fig. 2d). Differences between the LAs are mainly located at the loops in between the β strands and the N- and C-termini (Extended Data Fig. 2e). One of the flat surfaces contains the calcium binding site that is rich in negatively charged residues (NC surface). By fitting the structure of LA3^26^ into the additional densities at the three sites, we could establish the orientation and contact interface of the bound LAs. The bound LAs at the four A sites in the asymmetric unit have two major different orientations, of which one has a rotation of ∼ 56° relative to the other around the calcium binding site (Fig. 1a-d). The two different orientations of the bound LAs suggests that the binding of probably different LAs at different A sites. In both orientations, the bound LAs have the NC surface buried in the cleft. Residue K156 of E2 is in close proximity to the negatively charged residues around the calcium binding site of the fitted LAs and could be the key residue in contact at site A (Fig. 1e). Fitting at site B showed that the bound LA is in contact with the β-ribbon connector and the N-terminal helix in the domain A of E2 (Figs. 1b-c, 1e). Similar to the LA at site A, the NC surface is buried at the interface. Residue K231 of the β-ribbon connector is in close proximity to the calcium binding site and could be the key residue that mediates the interactions. Given the connecting densities between sites A and B, the bound LAs on sites A and B could be two consecutive LA repeats. Fitting at site C showed that the bound LA has close contacts with the E2 B domain, again with the NC surface buried at the interface (Fig. 1e). Similarly, a positively charged residue, K206 of E2, is in close proximity to the negatively charged residues around the calcium binding site and could be the key residue that mediates the interactions (Fig. 1e).

## Mapping of the essential LAs for binding EEEV PE6

To investigate the binding of each single LA to EEEV PE6, we coated single LAs on the probe and measured their direct interactions with the EEEV PE6 VLP by using biolayer interferometry (BLI). BLI is highly sensitive to the thickness change upon the attachment of the ligand^31^. Binding of EEEV VLP on the probe produce significant signals due to its big size. In addition, the multivalent interactions between the dense LA molecules on the probe and the EEEV VLP allow the detection of weak binding by a single LA (Extended Data Fig. 3a). Thus, this method can be used for qualitative detection of weak interactions between each single LA and EEEV VLP^16^. The BLI results showed that LA1-Fc, LA2-Fc, LA3-Fc, LA5-Fc and LA6-Fc can bind the EEEV PE6 VLP, whereas LA4-Fc, LA7-Fc and LA8-Fc do not have detectable signal of the bound EEEV PE6 VLP (Extended Data Fig. 3a).

Based on the analyses with the fitted LAs, we further mutated the potential key residues on the VLP and produced the mutant EEEV PE6-K156A, EEEV PE6-K206A and EEEV PE6-K231A VLPs, which are supposed to disrupt the binding at sites A, C and B, respectively. These mutant VLPs were then used for detecting site specific interactions with the LAs by BLI. The results showed that LA1-Fc and LA5-Fc can not bind the EEEV PE6-K156A VLP (Extended Data Fig. 3a), in which the binding site A in the cleft has been potentially disrupted. LA1-Fc, LA2-Fc, LA3-Fc, LA5-Fc and LA6- Fc can still bind the EEEV PE6-K206A and EEEV PE6-K231A VLPs, in which either the binding site C or B has been potentially disrupted (Extended Data Fig.3a). Additionally, we produced the E2 domain B and detected the interactions of E2 domain B with individual single LAs by using biolayer interferometry (BLI). The results showed that none of the single LAs can bind the E2 domain B with a detectable signal, indicating that single LA may only have undetectable weak interactions with site C on the E2 domain B (Extended Data Fig. 3a). LA1-8-Fc binds the E2 domain B with a KD of only 1.4 μM (Extended Data Fig. 3a), suggesting that multiple consecutive LAs can establish stable weak interactions with the E2 domain B or site C.

To further investigate the site preference of LA2, LA3 and LA6, we produced the mutant EEEV PE6-K156A-K231A, EEEV PE6-K206A-K231A and EEEV PE6-K156A-K206A VLPs, of which only one functional binding site was kept. BLI analysis with the VLPs carrying the double mutations showed that none of the LAs binds EEEV PE6-K156A-K206A (Extended Data Fig. 3a). Since EEEV PE6-K156A-K206A has, presumably, only site B for binding the receptor, indicating that binding to the site B may require the synergistic binding of the neighboring site A. In addition, the results also imply that for single LAs, no additional detectable binding site exists except for the three sites observed in the reconstruction. Furthermore, the results from EEEV PE6- K156A-K206A confirmed the essential role of K156 and K206 at site A and site C, respectively. The results with EEEV PE6-K156A-K231A are similar to these obtained with EEEV PE6-K156A (Extended Data Fig. 3a). LA1, LA2, LA3, LA5 and LA6 all can bind EEEV PE6-K206A-K231A (Extended Data Fig. 3a), of which only site A is functional. These data combined indicate that LA1 and LA5 specifically bind site A, whereas LA2, LA3 and LA6 can bind both sites A and C and binding of site B probably requires more than one LA (Fig. 2a).

**Fig. 2.**
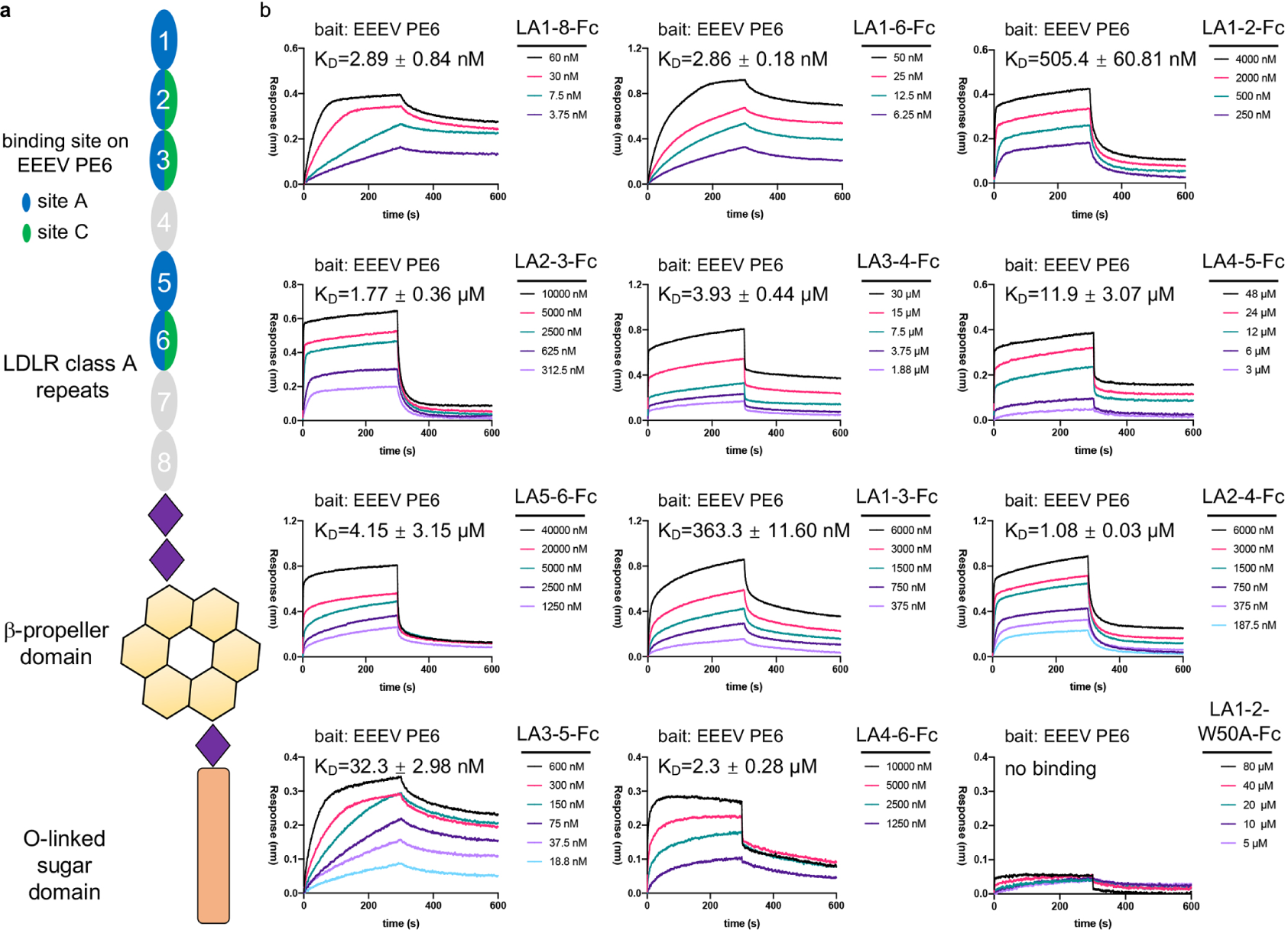
BLI analysis of the binding of different 2LAs and 3LAs to the EEEV PE6 VLP. **a**, A schematic diagram showing the ectodomains of VLDLR. The LA repeats are colored according to their binding specificity to the three sites on the E spike of EEEV PE6. **b**, BLI analysis showing the binding of VLDLR fragments that contains either 2 consecutive LAs (2LA) or three consecutive LAs (3LA) to the EEEV PE6 VLP. Data from one representative measurement of two independent experiments are presented. For each sample, the KD was calculated by averaging the values from the two independent experiments.

## Interactions between consecutive LA repeats and EEEV PE6

The BLI data with single LAs suggested that the binding at site B may involve more than one LA, most likely two consecutive LAs that occupy both the sites A and B. We further investigated the interactions between consecutive LA repeats and the EEEV PE6 VLP by using BLI. Constructs that contain LA7 or LA8 were not made since these LA repeats showed no detectable interaction with the EEEV PE6 VLP (Extended Data Fig.3a) and are located at the membrane-proximate end of the receptor. To measure the binding kinetic parameters precisely, we coated the anti-EEEV-E2 antibody EEEV-143^13^ on the probe. Then the VLPs were coated on the probe through the antibody and used for detecting the interactions with the LA repeats. By using this method, the binding of LA1-8 to the EEEV PE6 VLP was measured and the results indicate a strong binding with a KD of 2.9 nM (Fig. 2b).

We made constructs that have two consecutive LAs (2LA constructs) and then measured their interactions with the EEEV PE6 VLP. The results showed that the 2LAs, including LA1-2-Fc (KD: 505.4 nM), LA2-3-Fc (KD: 1.8 μM), LA3-4-Fc (KD: 3.9 μM), LA4-5-Fc (KD: 11.9 μM) and LA5-6-Fc (KD: 4.2 μM), have only weak interactions with the EEEV PE6 VLP (Fig. 2b). Among these, LA1-2-Fc has a relatively higher binding affinity.

Site C is far away from either site A or site B. At least three or four consecutive LAs would be required for simultaneous occupy of sites A and C or sites B and C. We next measured the interactions between fragments that have three consecutive LAs (3LAs) and the EEEV PE6 VLP. The results showed that LA3-5-Fc can bind the VLP much stronger compared with other 3LAs, including LA1-3-Fc, LA2-4-Fc, and LA4- 6-Fc. The measured KD of LA3-5-Fc is 32.3 nM, which is one to two orders of magnitude smaller than these of LA1-3-Fc (KD: 363.3 nM), LA2-4-Fc (KD: 1.1 μM) and LA4-6-Fc (KD: 2.3 μM) (Fig. 2b). The dramatic increase in the binding of LA3-5- Fc, but not other 3LA constructs, suggest possible synergistic binding through the LAs of LA3-5. Addition of a single LA repeat around LA3-5 did not further improve the binding affinity, as indicated by the measured KD values of LA3-6-Fc (KD: 71.5 nM) and LA2-5-Fc (KD: 42.5 nM) (Extended Data Fig. 3b). However, further addition of the LA repeat at either the N-terminus of LA2-5 (LA1-5-Fc) or the C-terminus of LA2-5 (LA2-6-Fc) improved the binding affinity to a level comparable to that of LA1-8-Fc (Extended Data Fig. 3b), indicating that compared with LA3-5, the introduction of two more additional LAs may create new binding modes.

## Structures of the VLDLR fragments in complex with EEEV PE6 VLP

The higher binding affinities of LA1-2-Fc and LA3-5-Fc suggest that these fragments may mediate different binding modes of the receptor. To further investigate the detailed binding modes of the LA repeats to EEEV PE6, we selected LA1-2-Fc and LA3-5-Fc for cryo-EM structural studies (Extended Data Figs. 4, 5, Extended Data Table 2).

The reconstruction of LA1-2-Fc shows strong additional densities on sites A and B (Fig. 3a). By treating the E proteins in one asymmetric unit of the icosahedral shell as a block, the reconstruction of the LA1-2-Fc and EEEV PE6 VLP complex was calculated to a resolution of 3.4 Å (Extended Data Fig. 4). Based on the density map, we identified LA1 at site A and LA2 at site B (Extended Data Fig. 4d). The structure model of LA1 could be built based on the density map (Extended Data Fig. 4d). Densities for the bound LA2 at site B are relatively weaker compared with these at site A. However, the key residues at the contact interface could be clearly identified (Extended Data Fig. 4d). LA1 is partially buried in the cleft created by two neighboring E1-E2 heterodimers and the long axis of LA1 is oriented approximately 30 degrees away from the 3 fold axis of the spike (Fig. 3a-c). LA1 interacts with the E proteins through mainly the two ridges (Fig. 3c). The negatively charged surface of LA1, that involves the calcium binding site and is located near one ridge, is buried and have direct contacts with the positively charged residue K156 of E2 (Figs. 3d, 3f). W50, which is in proximity to the calcium binding site of LA1, packs in parallel against the key residue E2-K156 (Fig. 3d). Previous studies showed that the tryptophan residue in proximity to the calcium binding site is critical for the binding of the LA repeat to SFV and VEEV^24,26^. To verify the role of W50, we made the mutant LA1-2-W50A-Fc and BLI analysis showed that W50A completely abolished the binding of LA1-2-Fc to the EEEV PE6 VLP (Fig. 2b). Residues K32, T41-R44 at the other ridge of LA1 have direct interactions with the fusion loop of E1 and residue H175 of E2, constituting the other major contact interface with the EEEV PE6 VLP (Fig. 3e). LA2 at the site B has only a small contact surface area of approximately 451 Å^2^ with the E proteins (Fig. 3g). The calcium binding site of LA2 is buried at the interface. K231 of E2 is pointing to the center of the calcium binding site, where D92, D94 and D96 of LA2 are in close proximity to K231 (Fig. 3g). To further verify the critic role of residues E2-K156 and E2-K231, we measured the binding of LA1-2-Fc to PE6-K156A and PE6-K231A. The BLI results showed that both mutations significantly affected the binding of LA1-2-Fc (Extended Data Fig. 3b).

**Fig. 3.**
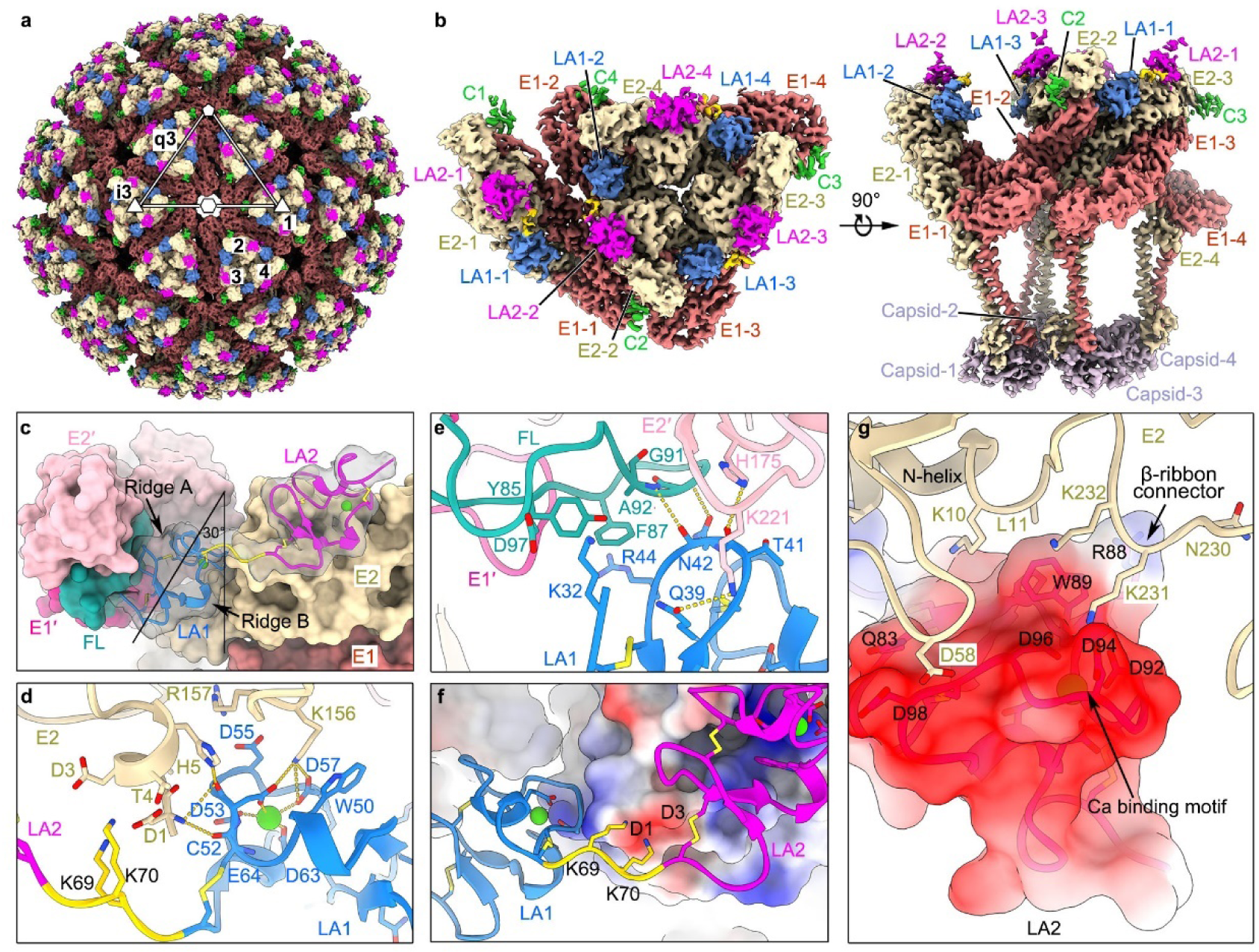
Structure of VLDLR LA1-2 in complex with the EEEV PE6 VLP. **a**, A surface rendered representation showing the density map of the EEEV PE6 VLP bound with LA1-2-Fc. The E1, E2, and capsid of the EEEV PE6 VLP are colored the same as in Fig. 1. Additional densities at three different locations on the spike, which could be contributed by the bound LA repeats, are colored blue, magenta and green, respectively. The labels i3 and q3 indicate one i3 spike and one q3 spike, respectively. **b**, A surface rendered representation showing the density map of four E1-E2 heterodimers in one asymmetric unit. The density map was calculated with the block- based reconstruction. The E proteins and the additional densities at three different sites on the spike are colored the same as in “a”. **c**, The bound VLDLR LA1-2 on the E1 and E2 glycoproteins of the EEEV PE6 VLP. The structures of VLDLR LA1-2 are shown in ribbon, and the structure of E proteins are shown in surface. The density map of VLDLR LA1-2, which is set transparent, was low pass filtered to a resolution of 6 Å and the contouring level was set to 0.00816 e^-^/ Å^3^. **d**,**e**, Detailed contacts between VLDLR LA1 and the EEEV PE6 E proteins at site A. **f**, Ribbon and surface-rendered representations showing contacts between VLDLR LA1-2 and the EEEV PE6 E proteins. The surface of the E proteins is colored according to the surface electrostatic potential. **g**, Ribbon and surface-rendered representations showing contacts between the VLDLR LA2 and the EEEV PE6 E2 at site B. The surface of LA2 is colored according to the surface electrostatic potential.

The reconstruction of the LA3-5-Fc and EEEV PE6 VLP complex was calculated to a resolution of 3.4 Å (Extended Data Fig. 5). The result shows additional densities only on sites A and C, indicating that the binding mode of LA3-5-Fc is distinctly different from that of LA1-2-Fc (Fig. 4). By fitting the structures of LA3 and LA5 into the additional densities, we could identify LA5 rather than LA3 at site A, based on the densities around residues F198 of LA5 and I118 of LA3, and V211 of LA5 and R133 of LA3 (Extended Data Fig. 5d), which are the residues at the equivalent positions of LA3 and LA5, respectively, but have distinct different features (Extended Data Fig. 5d). The results are consistent with the BLI analysis, which showed that LA5 is site A specific (Extended Data Fig. 3a). The atomic model of the bound LA5 could be built based on the density map (Extended Data Fig. 5e). LA5 binds site A mainly through only one ridge and the buried surface area at the interface is only 302 A^2^. The negatively charged residues around the bound calcium of LA5 are buried and have close contacts with residue K156 of E2 (Fig. 4d). Although the major area that LA1 and LA5 attach to is similar, LA5 binds site A in a distinctly different orientation when compared with that of LA1. The long axis of the bound LA5 is almost parallel to the axis of the trimeric spike (Fig. 4e). The structure of LA3 could be well fitted into the additional densities at site C and densities of the residues at the contact interface could be clearly observed (Extended Data Fig. 5f). At the contact interface, residues Q126, D135, E137 and D139 of LA3 form salt bridges and hydrogen bonds with residues of the E proteins, while W132 of LA3 is parallelly packed with residue K206 and V204 of E2 (Fig. 4f). Compared with other LAs, residue E137 of LA3 is unique (Extended Data Fig. 6a) and could help to establish additional salt bridges with K61 of E1 and K206 of E2 (Fig. 4f). Mutating E137 to the residue D137 as in other LAs, significantly reduced the binding affinity of LA3-5-Fc (Extended Data Fig. 3b). At a low contouring level, densities for LA4 could be observed in between the LA3 on one spike and the LA5 on a neighboring spike (Fig. 4c), indicating that LA3-5 crosslinks two neighboring spikes. Further mutagenesis studies showed that either W132G of the LA3 or D217A of LA5 abolishes the binding of LA3-5 to the EEEV PE6 VLP (Fig. 4g), indicating that W132 and D217 are the key residues of LA3 and LA5 for binding EEEV PE6, respectively. These results also confirm that binding of LA3-5 to EEEV PE6 requires the synergistic action of both LA3 and LA5. The binding of LA3-5 require the A site as does LA1-2, indicating that these fragments can compete each other for binding site A. To further verify the critic role of residue E2-K206, we measured the binding of LA3-5-Fc to PE6-K206A. The BLI results showed that the mutation K206A significantly affected the binding of LA3- 5-Fc (Extended Data Fig. 3b).

**Fig. 4.**
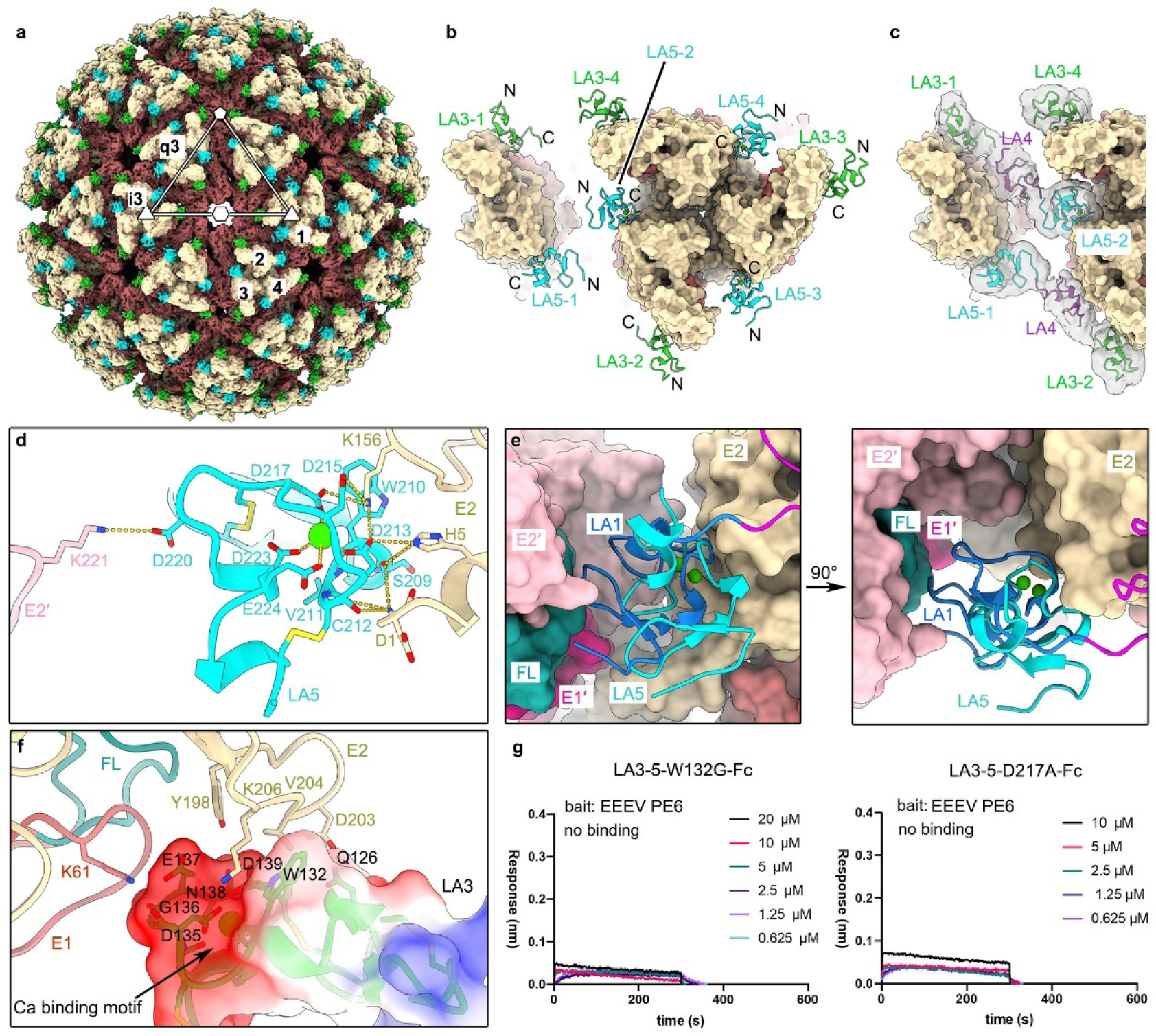
Structure of VLDLR LA3-5 in complex with the EEEV PE6 VLP. **a**, A surface rendered representation showing the overall structure of the complex. The E1, E2, and capsid of EEEV PE6 VLP are colored the same as in Fig. 1. Additional densities at two different locations on the sites A and C of the spike, which could be contributed by the bound LA repeats, are colored cyan and green, respectively. The labels i3 and q3 indicate one i3 spike and one q3 spike, respectively. **b**, Ribbon and surface rendered representations showing the density map of one asymmetric unit. The density map was obtained from the block-based reconstruction that treats one asymmetric unit as an individual block. The E proteins and the additional densities at two different sites on the spike are colored the same as in “a”. **c**, Ribbon and surface rendered representations showing the bound LAs on the spike. Additional densities beyond the spike, which were set transparent, were low pass filtered to a resolution of 8 Å and the contouring level was set to 0.0025 e^-^/ Å^3^. A model of LA4 predicted by AlphaFold was fitted in the densities with the N- and C-termini of LA4 posed toward the C-termini of LA3 at site C and N-termini of LA5 at site A, respectively. **d**, Detailed contacts between VLDLR LA5 and the EEEV PE6 E proteins at site A. **e**, Structural comparisons of the bound LA1 and LA5 at site A. **f**, Ribbon and surface-rendered representations showing contacts between VLDLR LA3 and the E proteins at site C. The surface of LA3 is colored according to the electrostatic potential. **g**, BLI analysis showing the binding of VLDLR LA3-5 mutants to the EEEV PE6 VLP.

## The multiple different binding modes of VLDLR

The combination of multiple bounders and binding sites can generate many different complex modes. The fragments LA3-5 and LA1-2 have much higher binding affinities compared with these with the same number of LA repeats. Additionally, the cryoEM reconstructions with LA1-2-Fc and LA3-5-Fc showed that the densities of LA3-5 and LA1-2 share similar features to these observed in the reconstruction of the LA1-8-Fc-PE6-VLP complex. The fragments LA1-2 and LA3-5 could play a major role in the binding of VLDLR to EEEV PE6. With the high resolution structures of LA1-2 and LA3-5, we further analyzed the reconstruction of the LA1-8-Fc and EEEV PE6 VLP complex, which shows strong densities at sites A and C, and weak densities at only sites B of the q3 spike. Among the three B sites on the q3 spike, the one (B3, Fig. 1a, b, d) near the 5 fold vertex has the strongest densities. The orientation of the densities at the nearby site A3 are consistent with these of the bound LA1 (Fig. 1a, b, d). In contrast, the orientation of the densities at the site A1 of the i3 spike are consistent with these of the bound LA5. Densities at site A2 and A4 are probably the average of the bound LAs. The fragment LA3-5 mediated binding is likely dominant, since the densities at sites B are much weaker compared to these on sites A and C.

It should be noted that the binding affinity of LA1-2 or LA3-5 is significantly weaker than that of LA1-8 or LA1-6, implying that, except for the LA1-2 and LA3-5 mediated binding modes, additional binding mode must exist when more than 4 consecutive LA repeats present. Sites C around the 2/5 fold axis may accommodate more than three consecutive LA repeats. To investigate possible additional binding modes with VLDLR fragments of more than 4 consecutive LA repeats, we incubated LA1-8-Fc and the EEEV PE6-K156A VLP, with which both the LA1-2 and LA3-5 mediated binding should be disabled. As expected, the cryoEM reconstruction of the complex showed additional densities only on sites C. Further structural analysis by fitting the structure of LA3 into the density map showed that the fragments of more than three consecutive LA repeats could bind the C sites around the 2/5 fold axis (Extended Data Fig. 6b).

## VLDLR mediated cell attachment of EEEV PE6

To further investigate whether the LA1-2 or LA3-5 mediated binding mode could establish stable attachment of the virus to the cell, we mutated the receptor and produced VLDLR-W132A and VLDLR-W50A, in which the key residue W132 of LA3 and W50 of LA1 were mutated to abolish the binding of LA3 and LA1, respectively. As a consequence, the LA3-5 or the LA1-2 mediated binding mode of VLDLR would be presumably disrupted. We built HEK293T cell lines that stably express either the wild type VLDLR, the LA3-dysfunctional (W132A) or the LA1-dysfunctional (W50A) VLDLR mutant. Cell attachment assays were then performed with the stable cell lines and the AF647 labeled EEEV PE6 VLP. As expected, the efficiency of the mutant VLDLR-W50A mediated cell attachment of the EEEV PE6 VLP was significantly reduced when compared with that of the wild type VLDLR mediated cell attachment (Fig. 5a-c). However, the efficiency of the mutant VLDLR-W132A mediated cell attachment was significantly enhanced when compared with that of the wild type VLDLR mediated cell attachment (Fig. 5a-c).

**Fig. 5.**
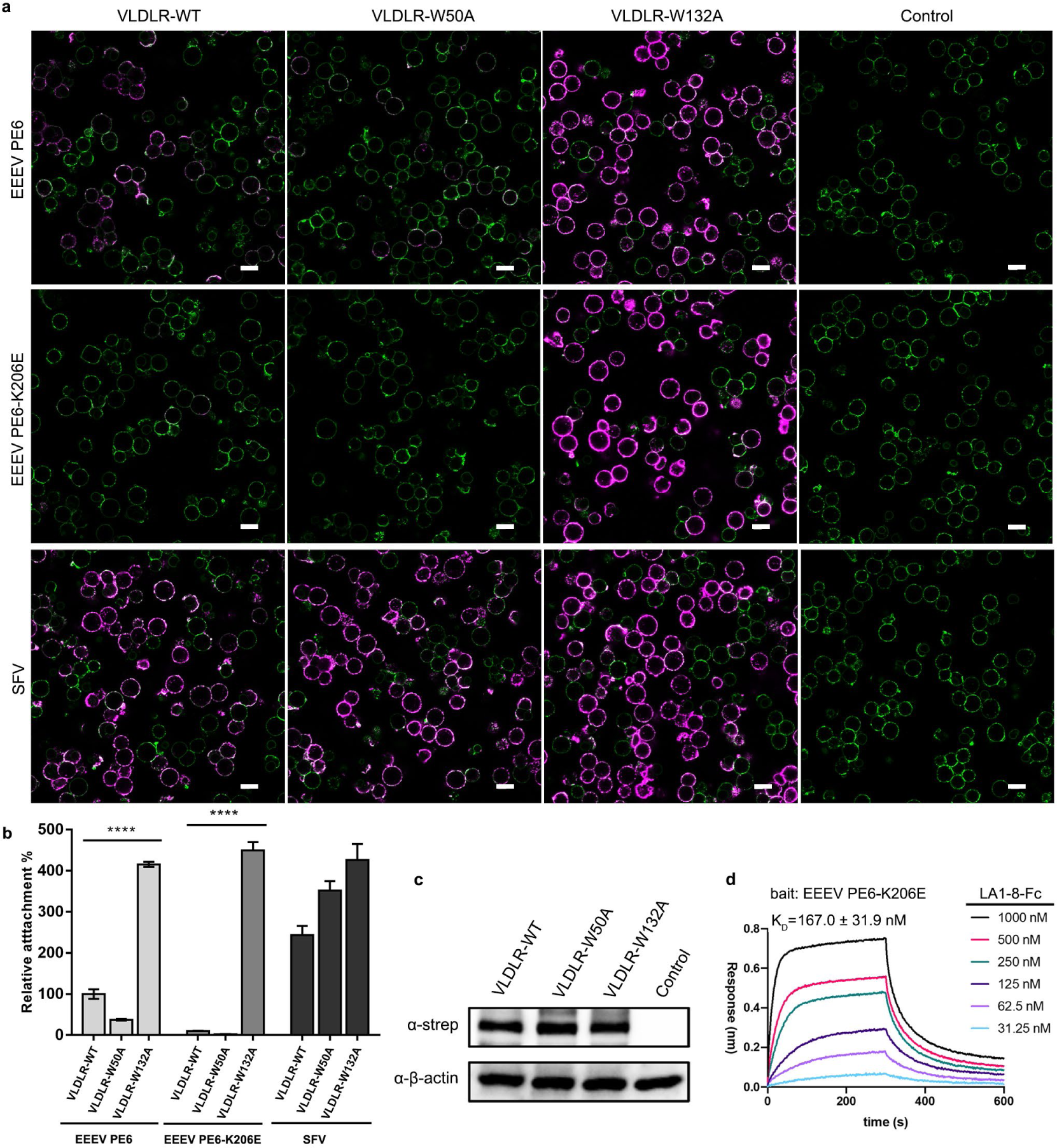
Cell attachment of VLDLR to different EEEV strains. **a**, Representative confocal microscopy images showing the attachment of EEEV to the HEK293T cell lines that express the wildtype VLDLR, VLDLR-W50A and VLDLR- W132A, respectively. The EEEV-PE6 and EEEV PE6-K206E VLPs were used. SFV VLPs were used as a control. Cells were incubated with the AF647 labeled VLPs at 37°C for 15 min and then were imaged by using a confocal microscopy. Fluorescence from the WGA-AF488 labeled cell membranes and the AF647 labeled VLPs were excited by laser beams with a wavelength of 488.6 nm for the wavelength of 639.8 nm, respectively. The power and PMT gain (HV) of the laser beam at the wavelength of 488.6 nm were set to 0.6 and 40 V, respectively. The power and PMT gain (HV) of the laser beam at the wavelength of 639.8 nm were set to 2 and 88 V, respectively. The scale bar is 20 μm. **b**, Relative attachment rate of the VLPs. Data are represented as mean ± SEM calculated with three groups of images from independent assays. Five images are in each group. One-way ANOVA with Tukey’s multiple comparisons test,****P<0.0001. **c**, Western blot analysis showing the expression level of the wildtype VLDLR, VLDLR-W50A and VLDLR-W132A in the same number of cells. **d**, BLI analysis showing the binding of VLDLR LA1-8-Fc to the EEEV PE6-K206E VLP. Data from one representative measurement of two independent experiments are presented. The KD was calculated by averaging the values from the two independent experiments.

We next explored the mechanism involved in the VLDLR-W132A mediated enhancement of EEEV cell attachment. We produced the mutant LA1-6-W132A-Fc. BLI analysis indicated that the mutation W132A does not affect the binding of LA1-6 (LA1-6-W132A-Fc, KD: 2.6 nM) (Extended Data Fig. 3b), suggesting that the loss of function in LA3 could be compensated by other LAs. CryoEM reconstructions of the LA1-6-Fc-W132A and EEEV PE6 VLP complex were then calculated (Fig. 6). The results show strong additional densities only on sites A and B and the densities on sites A and B are equally strong. The density distribution is similar to that of the LA1-2-Fc- PE6-VLP complex, indicating that the LA1-2 mediated binding to EEEV PE6 is dominant with the mutant LA1-6-Fc-W132A. Introduction of the dysfunctional LA3 in VLDLR drives the switch from the LA3-5 dominated binding mode to the LA1-2 dominated binding mode. LA3 is the determinant of the binding mode and may interfere with the binding of LA1-2. The conclusion could be further supported by the mutant LA1-3-W132A-Fc (KD: 279.0 nM) (Extended Data Fig. 3b), which has slightly higher binding affinity to the EEEV PE6 VLP compared with that of the wild type fragment LA1-3-Fc (KD: 363.3 nM) (Fig. 2b). Furthermore, we determined the cryoEM structure of LA1-3-Fc in complex with the EEEV PE6 VLP and checked the possible disturbance on the binding of LA1-2 by LA3. Compared with the reconstruction of the LA1-2-Fc and EEEV PE6 VLP complex, the densities at sites B were largely eliminated in the reconstruction of the LA1-3-Fc and EEEV PE6 VLP complex (Fig. 6), indicating that LA3 can compete with LA1 for binding site A and disturbs the binding of LA1-2. Similarly, we did the cryoEM structural studies of the LA2-3-Fc-PE6-VLP complex (Fig. 6). We observed additional densities only at sites A. Comparison of the density maps showed that the bound LA has a similar orientation as that of LA5 (Extended Data Fig. 6c). Given that both LA2 and LA3 can bind site A, the densities at sites A could be the average of LA2 and LA3. No matter LA2 or LA3 binds to site A, the other LA is highly flexible and not visible, which is quite different when compared with the binding of LA1-2.

**Fig. 6.**
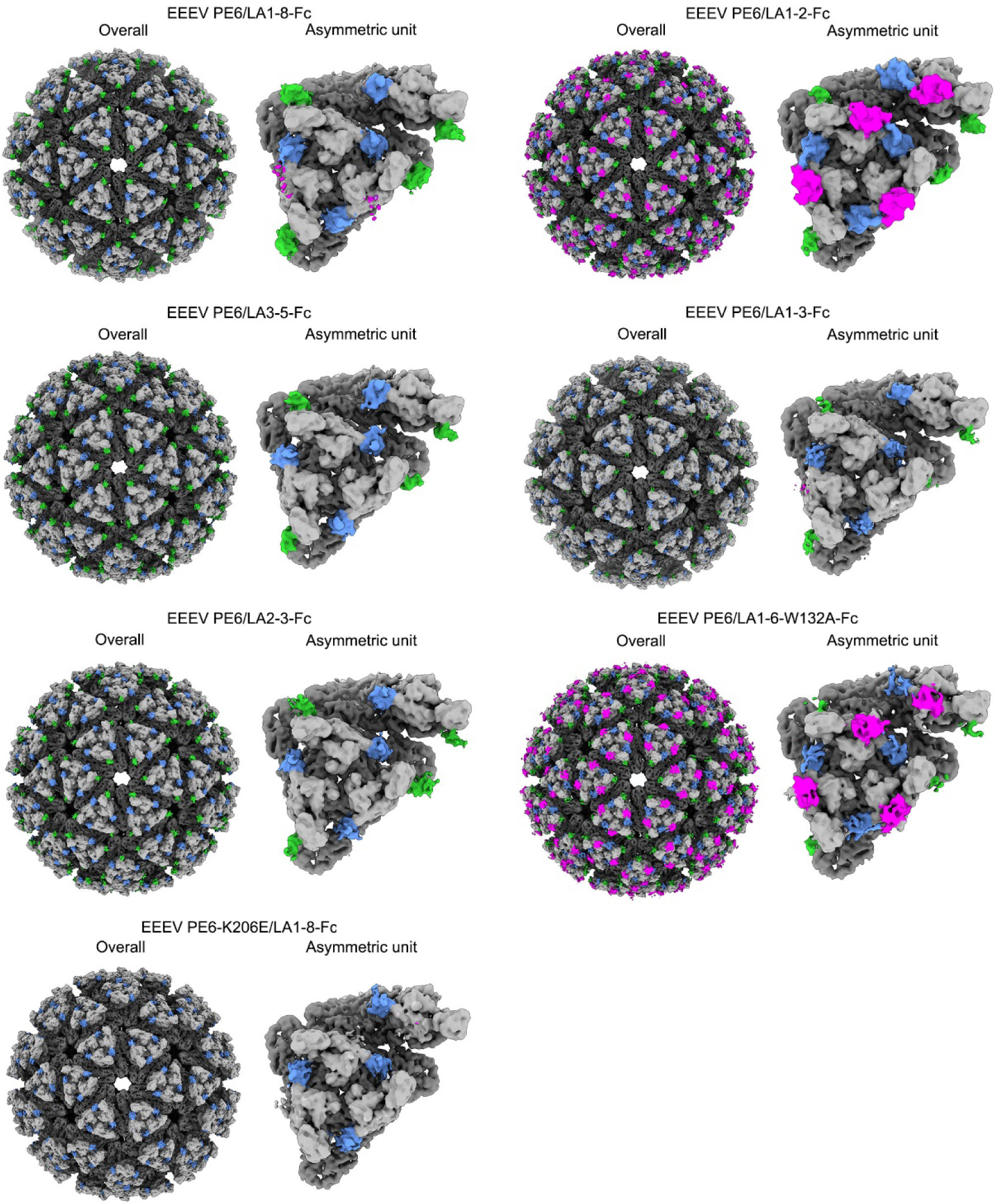
Comparison of different binding modes of VLDLR to the EEEV VLP. Surface-rendered representations showing the overall structures and the asymmetric unit of the EEEV PE6 VLP in complex with VLDLR LA1-8-Fc, LA1-2-Fc, LA3-5-Fc, LA1-3-Fc, LA2-3-Fc, LA1-6-W132A-Fc, and the EEEV PE6-K206E VLP in complex with LA 1-8-Fc. The densities of E1 and E2 proteins are colored dark gray and light gray, respectively. The additional densities at sites A, B and C are colored blue, magenta and green, respectively. The density maps of the asymmetric unit were low passed to 6 Å and normalized with the densities of the selected region of E2 (residues 80-120), which are located at the central region of the spike.

Taking together, these data combined indicate that LA3 can compete with LA1 for binding site A and thus, disturbs the LA1-2 mediated stable interactions with the virion. Both the LA1-2 and the LA3-5 mediated interactions could establish stable attachment of the virion to the cell, but the LA1-2 mediated attachment may mediate virus attachment more efficiently, due to probably the defined orientation of the bound receptor created by LA1-2. The C sites around the 5/2-fold symmetry axis generate additional binding modes of VLDLR, which could also play an important role in mediating the virus entry.

## Binding of VLDLR to different EEEV strains

Of note, sequence alignments indicate that the envelope proteins of the PE6 strain have two unique mutations when compared with these of other EEEV strains, including the North and South American strains (Extended Data Figs. 7, 8). One of the unique mutations is the residue K206 of E2, which is E in other virulent strains. K206 is the key residue of site C, while the residue E206 may abolish or significantly weaken the binding of VLDLR at site C. We, thus, mutated K206 to E206 and made the mutant EEEV PE6-K206E VLP. BLI analysis showed that the binding of LA1-8-Fc was significantly affected by the mutation K206E. LA1-8-Fc binds to the EEEV PE6- K206E VLP with a KD of 167.0 nM (Fig. 5d). In contrast, the KD with the EEEV PE6 VLP is 2.9 nM (Fig. 2b). Further cryoEM structural analysis of LA1-8-Fc in complex with the EEEV PE6-K206E VLP showed that similar additional densities on sites A and B as these of the LA1-8-Fc-PE6-VLP complex (Fig. 6). However, no additional densities could be observed on sites C, indicating that the C sites are impaired by the mutation K206E.

We next measured the VLDLR mediated cell attachment with the EEEV PE6- K206E VLP. The results showed that the mutation K206E significantly reduced the cell attachment of EEEV (Fig. 5a-c and Extended Data Fig. 9), which is in consistent with the BLI analysis (Figs. 2b, 5d). We further measured the VLDLR-W132A mediated cell attachment of the EEEV PE6-K206E VLP. The results showed that the mutation W132A promoted approximately 40 times more VLPs attached to the cells when compared with that of the wild type VLDLR mediated cell attachment (Fig. 5a-c and Extended Data Fig. 9).

## Discussion

### Selective binding of the LAs at different sites

Sequence alignments show that the key residues involved in binding VLDLR at sites A and C are highly conserved among LA1, LA2, LA3, LA5 and LA6 (Extended Data Fig. 6a). Comparing to the LAs that can bind site C, LA1 and LA5 have different residues at positions 44 and 47 of LA1 and positions 204 and 214 of LA5. Mutagenesis studies showed that LA1-R44Q-T47P-Fc and LA5-S204Q-D214G-Fc can bind EEEV PE6-K156A (Extended Data Fig. 10a), indicating these are the key residues that prevent the binding of LA1 and LA5 at site C. Structural superimpositions showed that LA2 and LA3 could bind site A in a similar way as that of LA5 (Extended Data Fig. 6c).

The bound LA1 places its C terminus towards site B, facilitates the binding of LA2 at site B, and promotes specific strong binding of LA1-2 at sites A and B, which distinguishes LA1-2 from other 2LAs (Figs. 2b, 3c). In addition, LA1-2 defines a unique orientation of the receptor that could favor the binding of cofactors on the cell membrane and help to build the stable attachment of the virus.

Site A and site C are 60 Å away from each other and they can only be simultaneously occupied by two LAs of a VLDLR fragment that contains at least three LAs. Location of the N and C-termini of the bound LAs further confines the binding of a N-terminal LA at site A from one spike and the binding of a C-terminal LA at site C from a neighboring spike. Among the 3LAs, only LA3-5 can satisfy the requirements for binding sites A and C at the same time, since LA1 is site A specific, while LA4 can not binding either site A or site C.

In conclusion, the difference in the binding affinity and specificities of each LA and the geometric restrains created by the arrangement of the viral glycoproteins, define the fragment mediated distinct binding modes of VLDLR to EEEV.

### The additional receptor binding site on the EEEV PE6 strain

The EEEV PE6 strain was isolated from the brain of an infected horse and was propagated in chicken embryos^32,33^. The E206K mutation of E2 in EEEV PE6 was considered as the key factor of attenuation since domain B was identified as the epitope of neutralization antibodies^29^. However, our results showed that the E206K mutation creates an additional binding site of the receptor and significantly enhanced the VLDLR mediated cell attachment of EEEV (Fig. 5a-c, Extended Data Fig. 6b). The other unique mutation identified in the EEEV PE6 envelop protein, F311 of E1, is not relevant to the binding of VLDLR (Extended Data Fig. 7). The key mutation that causes the attenuation of EEEV PE6 could be in the none structural proteins rather than the structural proteins.

There are advantages and disadvantages of using the EEEV PE6 strain as vaccines. In its inactivated or VLP form, EEEV PE6 strain could provide protection against the additional C site at the domain B of E2, which could exist in the circulating enzootic EEEV strains. However, if being used as the live attenuated vaccine, EEEV PE6 strain can have the risk of being converted into a strain of strong virulence, due to its high efficiency in binding VLDLR.

### Enhanced cell attachment caused by the dysfunctional LA3

Of note, we observed that the VLDLR mediated cell attachment of EEEV was significantly enhanced with the mutation W132A, which results in a dysfunctional LA3. The enhancement was observed for both the EEEV PE6 and PE6-K206E VLPs (Fig. 5a-c, Extended Data Fig. 9) and the promotion on PE6-K206E is dramatical. Considering that E206 is dominant in the current virulent EEEV strains, our finding might be of physiological importance. The mutation W132G, which is supposed to behave similarly as W132A, can be identified in human genome and SNP sequences. W132G was shown to abolish the binding of LA3-5-Fc to the EEEV PE6 VLP (Fig. 4g). We further made the mutant VLDLR-W132G and the cell attachment assays showed similar results as these with the mutant VLDLR-W132A (Extended Data Fig. 10b). It would be interesting to investigate in the future whether human carrying the W132G mutation in VLDLR or other similar mutations in LA3 are more susceptible and of higher risk to EEEV infection.

### VLDLR mediated infection of EEEV across different species

Previous studies showed that the VLDLR from *S. vulgaris*, the reservoir host of EEEV, can not mediated the infection of K562 cells by EEEV^16^. However, sequence alignments of the VLDLRs from different species showed that key residues of the *S. vulgaris* VLDLR are exactly the same as these of the human VLDLR (Extended Data Fig. 11). We further measured the binding of *S. vulgaris* VLDLR LA1-8-Fc (LA1-8- Fc-*S. vulgaris*) to the EEEV PE6 VLP and the results showed that LA1-8-Fc-*S. vulgaris* binds the EEEV PE6 VLP with a KD of 11.0 nM, which is similar as that of the human LA1-8-Fc (Extended Data Fig. 12a and Fig. 2b).

We then investigated the location of the overexpressed *S. vulgaris* VLDLR in HEK293T cells and the results showed that the distribution of the receptor has no significant difference when compared with that of the human VLDLR (Extended Data Fig. 12b). Next, we checked the attachment of the EEEV PE6 VLP to the HEK293T cells that overexpress the *S. vulgaris* VLDLR and the results showed that *S. vulgaris* VLDLR can mediate the stable attachment of the EEEV PE6 VLP. Furthermore, the VLDLR mediated entry of the EEEV PE6 VLP was checked and the results showed the labeled EEEV PE6 VLP particles in the cell (Extended Data Fig. 12c). The binding of EEEV PE6-K206E to the *S. vulgaris* VLDLR^+^ HEK293T cells is weak but can be detected (Extended Data Fig. 12d). These results indicate that the *S. vulgaris* VLDLR is able to mediate the attachment and entry of EEEV. However, different EEEV strains may have different susceptibility to the bird hosts.

### VLDLR mediated infection that cross different species

Sequence alignments of VLDLRs from the vertebrate species show that the key residues involved in the interactions with sites A, B and C are completely conserved in most of the LA repeats (Extended Data Fig. 11). Thus, the LA1-2 and LA3-5 mediated binding of VLDLR to EEEV could be well maintained in most vertebrate species. Larger variations are observed in the sequences of VLDLR from invertebrate species when compared with these among the vertebrate species (Extended Data Fig. 11). However, residues involved in direct contacts with EEEV are still largely conserved in the invertebrate VLDLRs (Extended Data Fig. 11), suggesting that the binding modes of VLDLR could be well maintained for mediating the infection of invertebrate vectors. The different binding modes of VLDLR involve different LAs. Loss of any functional LA may only drive the switch between different binding modes and does not affect the attachment of EEEV to the cells. Thus, EEEV could be tolerant to the variations in VLDLRs from different species. The mechanism involved in the tolerance of EEEV to the variation of VLDLR is quite different from that of SFV. LA1-5 of VLDLR binds multiple E1-DIII sites of SFV through a synergistic mode, in which the binding affinity of a single LA could be sacrificed for maximizing the overall binding affinity built on multiple sites^26^.

### Implication for the VLDLR mediated infection of other viruses

Sequence alignments of alphavirus E proteins show that the key residues involved in binding VLDLR is not conserved among E proteins of alphaviruses (Extended Data Fig. 13a). Thus, it is unlikely that VLDLR can mediate the entry of other alphaviruses in a similar way as does for EEEV.

Comparisons of the binding of the LA repeat to different viruses, the binding modes are quite different even at the similar site, such as the cleft in between the E1-E2 heterodimers. The D1 domain of LDLRAD3 binds VEEV at a deeper position of the cleft, with significant more contacts, and much higher affinity when compared with the binding of LA1 or LA5 to EEEV (Extended Data Fig. 13b). However, a common feature is that a positively charged residue of the virus surface protein interacts with the negatively charged residues around the calcium binding site of the LA repeat. The surface in contact could be as small as 245 Å^2^, such as these between LA3 and the E2 B domain of EEEV. The LA repeat could be a modular scaffold for specific binding of many different molecules, through either a single LA repeat or multiple LA repeats. In these involved multiple similar repeating subunits, although the binding affinity of each single LA is weak, simultaneous interactions at multiple sites could build up enough affinity for highly specific recognition of the target molecules, especially for these densely arranged molecules, such as viral envelop proteins and capsids.

## References

1 Lindsey, N. P., Martin, S. W., Staples, J. E. & Fischer, M. Notes from the Field: Multistate Outbreak of Eastern Equine Encephalitis Virus - United States, 2019. MMWR Morb Mortal Wkly Rep 69, 50-51, doi:10.15585/mmwr.mm6902a4 (2020).

2 de Novaes Oliveira, R., et al. Eastern equine encephalitis cases among horses in Brazil between 2005 and 2009. Arch. Virol. 159, 2615–2620, doi:10.1007/s00705-014-2121-4 (2014).

3 Lindsey, N. P., Staples, J. E. & Fischer, M. Eastern Equine Encephalitis Virus in the United States, 2003–2016. The American Journal of Tropical Medicine and Hygiene 98, 1472-1477, doi:10.4269/ajtmh.17-0927 (2018).

4 Armstrong, P. M. & Andreadis, T. G. Eastern equine encephalitis virus in mosquitoes and their role as bridge vectors. Emerg Infect Dis 16, 1869–1874, doi:10.3201/eid1612.100640 (2010).

5 Sherwood, J. A., Stehman, S. V., Howard, J. J. & Oliver, J. Cases of Eastern equine encephalitis in humans associated with Aedes canadensis, Coquillettidia perturbans and Culiseta melanura mosquitoes with the virus in New York State from 1971 to 2012 by analysis of aggregated published data. Epidemiol. Infect. 148, e72, doi:10.1017/s0950268820000308 (2020).

6 Sah, R. et al. The emerging scenario for the Eastern equine encephalitis virus and mitigation strategies to counteract this deadly mosquito-borne zoonotic virus, the cause of the most severe arboviral encephalitis in humans—an update. Frontiers in Tropical Diseases 3, doi:10.3389/fitd.2022.1077962 (2023).

7 Aréchiga-Ceballos, N. & Aguilar-Setién, A. Alphaviral equine encephalomyelitis (Eastern, Western and Venezuelan). Rev Sci Tech 34, 491–501, doi:10.20506/rst.34.2.2374 (2015).

8 Arrigo, N. C., Adams, A. P. & Weaver, S. C. Evolutionary patterns of eastern equine encephalitis virus in North versus South America suggest ecological differences and taxonomic revision. J. Virol. 84, 1014–1025, doi:10.1128/jvi.01586-09 (2010).

9 Ciota, A. T. Eastern Equine Encephalitis Virus Taxonomy, Genomics, and Evolution. J. Med. Entomol. 59, 14–19, doi:10.1093/jme/tjab079 (2022).

10 Kim, A. S. & Diamond, M. S. A molecular understanding of alphavirus entry and antibody protection. Nature Reviews Microbiology 21, 396–407, doi:10.1038/s41579-022-00825-7 (2023).

11 Holmes, A. C., Basore, K., Fremont, D. H. & Diamond, M. S. A molecular understanding of alphavirus entry. PLoS pathog. 16, e1008876, doi:10.1371/journal.ppat.1008876 (2020).

12 Sutton, M. S. et al. Vaccine elicitation and structural basis for antibody protection against alphaviruses. Cell 186, 2672–2689.e2625, doi:10.1016/j.cell.2023.05.019 (2023).

13 Williamson, L. E. et al. Human Antibodies Protect against Aerosolized Eastern Equine Encephalitis Virus Infection. Cell 183, 1884–1900.e1823, doi:10.1016/j.cell.2020.11.011 (2020).

14 Chen, C.-L. et al. Cryo-EM structure of eastern equine encephalitis virus in complex with heparan sulfate analogues. Proceedings of the National Academy of Sciences 117, 8890–8899, doi:doi:10.1073/pnas.1910670117 (2020).

15 Hasan, S. S. et al. Cryo-EM Structures of Eastern Equine Encephalitis Virus Reveal Mechanisms of Virus Disassembly and Antibody Neutralization. Cell Rep 25, 3136–3147.e3135, doi:10.1016/j.celrep.2018.11.067 (2018).

16 Clark, L. E. et al. VLDLR and ApoER2 are receptors for multiple alphaviruses. Nature 602, 475–480, doi:10.1038/s41586-021-04326-0 (2022).

17 Ma, H. et al. LDLRAD3 is a receptor for Venezuelan equine encephalitis virus. Nature 588, 308–314, doi:10.1038/s41586-020-2915-3 (2020).

18 Zimmerman, O. et al. Vertebrate-class-specific binding modes of the alphavirus receptor MXRA8. Cell 186, 4818–4833.e4825, doi:10.1016/j.cell.2023.09.007 (2023).

19 Zhang, R. et al. Mxra8 is a receptor for multiple arthritogenic alphaviruses. Nature 557, 570–574, doi:10.1038/s41586-018-0121-3 (2018).

20 Cao, Y., Wang, H., Jin, P., Ma, F. & Zhou, X. Identification and Characterization of the Very-Low-Density Lipoprotein Receptor Gene from Branchiostoma belcheri: Insights into the Origin and Evolution of the Low- Density Lipoprotein Receptor Gene Family. Animals (Basel) 13, doi:10.3390/ani13132193 (2023).

21 Nimpf, J. & Schneider, W. J. From cholesterol transport to signal transduction: low density lipoprotein receptor, very low density lipoprotein receptor, and apolipoprotein E receptor-2. Biochim. Biophys. Acta Mol. Cell Biol. Lipids. 1529, 287–298, doi:10.1016/S1388-1981(00)00155-4 (2000).

22 Fass, D., Blacklow, S., Kim, P. S. & Berger, J. M. Molecular basis of familial hypercholesterolaemia from structure of LDL receptor module. Nature 388, 691–693, doi:10.1038/41798 (1997).

23 Verdaguer, N., Fita, I., Reithmayer, M., Moser, R. & Blaas, D. X-ray structure of a minor group human rhinovirus bound to a fragment of its cellular receptor protein. Nat. Struct. Mol. Biol. 11, 429–434, doi:10.1038/nsmb753 (2004).

24 Ma, B., Huang, C., Ma, J., Xiang, Y. & Zhang, X. Structure of Venezuelan equine encephalitis virus with its receptor LDLRAD3. Nature 598, 677–681, doi:10.1038/s41586-021-03909-1 (2021).

25 Basore, K. et al. Structure of Venezuelan equine encephalitis virus in complex with the LDLRAD3 receptor. Nature 598, 672–676, doi:10.1038/s41586-021-03963-9 (2021).

26 Cao, D., Ma, B., Cao, Z., Zhang, X. & Xiang, Y. Structure of Semliki Forest virus in complex with its receptor VLDLR. Cell 186, 2208–2218.e2215, doi:10.1016/j.cell.2023.03.032 (2023).

27 Song, H. et al. Molecular basis of arthritogenic alphavirus receptor MXRA8 binding to chikungunya virus envelope protein. Cell 177, 1714–1724. e1712, doi:10.1016/j.cell.2019.04.008 (2019).

28 Basore, K. et al. Cryo-EM structure of chikungunya virus in complex with the Mxra8 receptor. Cell 177, 1725–1737. e1716, doi:10.1016/j.cell.2019.04.006 (2019).

29 Platteborze, P. L., Kondig, J. P., Schoepp, R. J. & Wasieloski, L. P. Comparative sequence analysis of the eastern equine encephalitis virus pathogenic strains FL91-4679 and GA97 to other North American strains. DNA Seq 16, 308–320, doi:10.1080/10425170500136889 (2005).

30 Jumper, J. et al. Highly accurate protein structure prediction with AlphaFold. Nature 596, 583–589, doi:10.1038/s41586-021-03819-2 (2021).

31. 31 Octet® BLI Label-Free Detection Systems for Biomolecular Interactions Analysis, >https://www.sartorius.com/en/products/protein-analysis/octet-bli-detection/octet-label-free-detection-systems> (2023).

32 Maire, L. F., 3rd, McKinney, R. W. & Cole, F. E., Jr. An inactivated eastern equine encephalomyelitis vaccine propagated in chick-embryo cell culture. I. Production and testing. Am. J. Trop. Med. Hyg. 19, 119–122, doi:10.4269/ajtmh.1970.19.119 (1970).

33 Bartelloni, P. J., McKinney, R. W., Duffy, T. P. & Cole, F. E., Jr. An inactivated eastern equine encephalomyelitis vaccine propagated in chick-embryo cell culture. II. Clinical and serologic responses in man. Am. J. Trop. Med. Hyg. 19, 123–126, doi:10.4269/ajtmh.1970.19.123 (1970).

